# *Plasmodium vivax-like* genome sequences shed new insights into *Plasmodium vivax* biology and evolution

**DOI:** 10.1101/205302

**Authors:** Aude Gilabert, Thomas D. Otto, Gavin G. Rutledge, Blaise Franzon, Benjamin Ollomo, Céline Arnathau, Patrick Durand, Nancy D. Moukodoum, Alain-Prince Okouga, Barthélémy Ngoubangoye, Boris Makanga, Larson Boudenga, Christophe Paupy, François Renaud, Frank Prugnolle, Virginie Rougeron

**Affiliations:** MIVEGEC, IRD, CNRS, Univ. Montpellier, Montpellier, FRANCE; Wellcome Trust Sanger Institute, Wellcome Trust Genome Campus, Cambridge CB10 1SA, UK; Institute of Infection, Immunity and Inflammation, University of Glasgow, College of Medical, Veterinary and Life Sciences, Sir Graeme Davies Building, 120 University Place, Glasgow G12 8TA, UK; Centre International de Recherches Médicales de Franceville, B.P. 769, Franceville, GABON

## Abstract

Although *Plasmodium vivax* is responsible for the majority of malaria infections outside Africa, little is known about its evolution and pathway to humans. Its closest genetic relative, *Plasmodium vivax-like*, was discovered in African great apes and is hypothesized to have given rise to *P. vivax* in humans. To unravel the evolutionary history and adaptation of *P. vivax*, we generated using long and short read sequence technologies the two first *P. vivax-like* reference genomes and 9 additional *P. vivax-like* genotypes. Analyses show that the genomes of *P. vivax* and *P. vivax-like* are highly similar and co-linear within the core regions. Phylogenetic analyses clearly show that *P. vivax-like* parasites form a genetically distinct clade from *P. vivax*. Concerning the relative divergence dating, we show that the evolution of *P. vivax* in humans did not occur at the same time as the other human malaria agents, thus suggesting that the transfer of *Plasmodium* parasites to humans happened several times independently over the history of the *Homo* genus. We further identify several key genes that exhibit signatures of positive selection exclusively in the human *P. vivax* parasites. Interestingly, two of these genes have been identified to also be under positive selection in the other main human malaria agent, *P. falciparum*, thus suggesting their key role in the evolution of the ability of these parasites to infect humans or their anthropophilic vectors. We finally demonstrate that some gene families important for red blood cell (RBC) invasion (a key step of the life cycle of these parasites) have undergone lineage-specific evolution in the human parasite (e.g. Reticulocyte Binding Proteins).

**Significance statements:** Among the five species responsible for this malaria in humans, *Plasmodium vivax* is the most prevalent outside Africa and causes severe and incapacitating clinical symptoms with significant effects on human health. Its closest known relative was recently discovered in African great apes, *Plasmodium vivax-like.* This study aims to characterize the genome of the closest ape-relative to the human *P. vivax* parasite in order to get a better understanding of the evolution of this parasite.

A total of eleven *P. vivax-like* samples were obtained from infected chimpanzee blood samples and an infected *Anopheles* mosquito collected in Gabon. Through technical accomplishment and using short and long read sequence technologies, two newly genomes of *P. vivax-like* and further nine additional draft sequences were obtained. The genome-wide analyses performed provided new insights into the biology and adaptive evolution of *P. vivax* to different host species.

## Main text

*Plasmodium vivax* is responsible for severe and incapacitating clinical symptoms in humans^1^. Traditionally, *P. vivax* has been neglected because of its lower mortality in comparison to *Plasmodium falciparum*^2,3^. Its ability to produce a dormant liver-stage form (hypnozoite), responsible for relapsing infections, makes it a challenging public health issue for malaria elimination. The recent emergence of antimalarial drug resistance^4^ as well as the discovery of severe and even fatal human cases^2,5,6^ has renewed the interest in this enigmatic species, including its evolutionary history and its origin in humans.

Earlier studies placed the origin of *P. vivax* in humans in Southeast Asia (“Out of Asia” hypothesis) based on its phylogenetic position in a clade of parasites infecting Asian monkeys^7^. At that time, the closest known relative of *P. vivax* was considered to be *Plasmodium cynomolgi*, an Asian monkey parasite^8^. However, this hypothesis was recently challenged with the discovery of another *Plasmodium* species, genetically closer to *P. vivax* than *P. cynomolgi*, circulating in African great apes (chimpanzees and gorillas)^9,10^. This new lineage (hereafter named *Plasmodium vivax-like*) was considered to have given rise to *P. vivax* in humans following a transfer of parasites from African apes^10^. But this “Out of Africa” hypothesis still remains debated. Moreover, a spillover of *P. vivax-like* parasites to human has been documented, thus making possible the release of new strains in new hosts species, specifically in human populations^9^.

In such a context, it seemed fundamental to characterize the genome of the closest ape-relative to the human *P. vivax* parasite in order to get a better understanding of the evolution of this parasite and also to identify the key genetic changes explaining the emergence of *P. vivax* in human populations.

### Genome Assemblies

Eleven *P. vivax-like* genotypes were obtained from two different kinds of samples: ten infected chimpanzee blood samples collected during successive routine sanitary controls of chimpanzees living in the Park of La Lékédi (a sanctuary in Gabon) and one infected *Anopheles* mosquito collected during an entomological survey carried out in the same park^11^ (Supplementary Table 1). For blood samples, white blood cells were depleted using the CF11 method^12^ to reduce the amount of host DNA. After DNA extraction, samples were subjected to whole genome amplification (WGA) in order to obtain sufficient parasite DNA for library preparation. Sequencing was then performed using short read Illumina technology. For one sample (Pvl06), long read sequencing (PacBio technology) was performed in order to get a better coverage of regions containing subtelomeric gene families.

**Table 1.** Genome features of the *P. vivax-like* Pvl01 (Illumina HiSeq sequenced) and Pvl06 (PacBio sequenced) strains, *P. vivax* reference strains SalI and PvP01^15^, *P. cynomolgi* B and M isolates^8,50^ and *P. knowlesi* H strain^17^. *Unassigned contigs indicated in parentheses.

|  | *P. vivax-like*<br>(Pvl01) | *P. vivax-like*<br>(Pvl06) | *P. vivax*<br>(PvP01) | *P. vivax*<br>(Sall) | *P. cynomolgi*<br>(B strain) | *P. cynomolgi*<br>(M strain) | *P. knowlesi*<br>(H strain) |
| --- | --- | --- | --- | --- | --- | --- | --- |
| **Assembly size (Mb)** | 27.5 | 18.8 | 29 | 26.8 | 26.2 | 30.6 | 24.1 |
| **Scaffolds** | 14 (226)* | 14 (370)* | 14 | 14 | 14 | 14 | 14 |
| **Overall G+C content (%)** | 44.9 | 45.8 | 39.8 | 42.3 | 40.340.4 | 37.3 | 37.5 |
| **Number of genes** | 5,532 | 4,953 | 6,642 | 6,690 | 5,722 | 6,632 | 5,188 |
| **Gene density (gene/Mb)** | 201.2 | 264.5 | 229 | 249.6 | 218.4 | 216.7 | 215.3 |
| **# 1:1 orthologues with PvP01** | 4,570 | 2,925 | - | 5,178 | 4,870 | 5,222 | 4,804 |

Among the eleven samples, ten presented mixed infections with other *Plasmodium* species (Supplementary Table 1). Four samples containing *P. gaboni* or *P. malariae-like* co-infections were used in other studies (see Supplementary Table 1)^13,14^. In order to obtain the *P. vivax-like* genotypes, sequencing reads were extracted based on their similarity to the reference genome sequence of *P. vivax*, PvP01^15^. Sequencing reads from two samples, one obtained using Illumina sequencing, Pvl01, and another using PacBio technology, Pvl06, were used to perform *de novo* genome assemblies and annotated to produce reference genomes for *P. vivax-like* (Supplementary Table 2). Of the two assemblies, Pvl01 is of considerably higher quality (4,570 orthologues to the PvP01 reference genome compared to 2,925 for Pvl06 (Table 1)). Both assemblies consist of 14 supercontigs (corresponding to the 14 *P. vivax* chromosomes) and, respectively for Pvl01 and Pvl06, 226 and 370 unassigned contigs, comprising a total of 27.5Mb and 18.8Mb in size respectively for Pvl01 and Pvl06. After annotation with Companion^16^, these two genomes contained 5,532 and 4,953 annotated genes (Table 1). The genome sequences obtained from the other samples were used for SNP calling, and population genetic and phylogenetic analyses.

**Table 2.** Multigene families copy number description of *P. vivax-like* (Pvl01 and Pvl06), *P. vivax* strains SalI and PvP01^15^, *P. cynomolgi* B and M strains^8,50^ and *P. knowlesi* H strain^17^. *For *P. vivax-like* Pvl01 and Pvl06, a non-exhaustive list of family genes is represented since only partial genomes were obtained. Pseudogeneized genes are included.

|  | *P. vivax-like*<br>(Pvl01)* | *P. vivax-like*<br>(Pvl06)* | *P. vivax*<br>(PvP01) | *P. vivax*<br>(Sall) | *P. cynomolgi*<br>(B strain) | *P. cynomolgi*<br>(M strain) | *P. knowlesi*<br>(H strain) |
| --- | --- | --- | --- | --- | --- | --- | --- |
| Vir/Pir<br>(Subtelomeric) | 148 | 14 | 1212 | 303 | 265 | 1373 | 64 |
| Msp3<br>(Central) | 9 | 0 | 12 | 11 | 12 | 14 | 3 |
| Msp7<br>(Central) | 11 | 12 | 13 | 13 | 13 | 11 | 5 |
| DBP<br>(Subtelomeric) | 1 | 0 | 2 | 1 | 2 | 2 | 3 |
| RBP<br>(Subtelomeric) | 9 | 3 | 10 | 9 | 8 | 6 | 2 |
| Pv-fam-a (trag)<br>(Subtelomeric) | 33 | 10 | 40 | 34 | 36 | 39 | 26 |
| Pv-fam-e (rad)<br>(Subtelomeric) | 38 | 15 | 40 | 34 | 27 | 27 | 16 |
| PST-A<br>(Subtelomeric<br>and central) | 6 | 3 | 10 | 11 | 9 | 8 | 7 |
| ETRAMP<br>(Subtelomeric) | 7 | 4 | 10 | 9 | 9 | 9 | 9 |
| CLAG (RhopH-1)<br>(Subtelomeric) | 4 | 2 | 3 | 3 | 2 | 2 | 2 |
| PvSTP1<br>(Subtelomeric) | 4 | 0 | 9 | 10 | 3 | 51 | 0 |
| PHIST (Pf-fam-b)<br>(Subtelomeric) | 20 | 12 | 84 | 64 | 48 | 54 | 15 |
| SERA<br>(Central) | 13 | 7 | 13 | 13 | 13 | 13 | 8 |

### Gene synteny and gene composition

Comparing the *P. vivax-like* reference genomes with those of *P. vivax* (PvP01 and SalI)^2,15^, *P. cynomolgi* (B strain)^8^ and *P. knowlesi* (H strain)^17^ reveals several similarities, including a similar GC content and extensive collinearity and conservation of gene content/organization (Table 1). The *P. vivax-like* core genome sequences are completely syntenic to the *P. vivax* PvP01 reference genome sequence (Supplementary Figures 1 and 2).

Because multigene families are known to evolve extremely rapidly in their genome structure, obtaining the full genomes of the closer species to the human *P. vivax* is fundamental to get a better understanding of its evolution, adaptation and emergence in different host species. For *Plasmodium* parasites, most species-specific genes are part of large gene families, such as *var* genes in *P. falciparum* or *pir* genes that currently are present in all *Plasmodium* genomes studied^18,19^. Table 2 provides a summarized view of gene content and copy number of the main multigene families in *P. vivax-like* in comparison to *P. vivax, P. knowlesi* and *P. cynomolgi*. Even if certain subtelomeric regions of our reference genomes (Supplementary Figures 1 and 2) are not complete, at least one copy of each major gene family was detected (Table 2). In comparison to *P. vivax* (as expected because of the partial subtelomeric sequencing coverage), the number of copies in each family was generally lower or equal in *P. vivax-like*. For these families, all genes were functional except for the CLAG (Cytoadherence-linked asexual gene) and SERA (Serine repeat antigen) families. For the CLAG family, all genes are functional except the one situated on chromosome 8 for *P. vivax-like* (confirmed for both Pvl01 and Pvl06) (Supplementary Figure 3). The CLAG family, strictly conserved in malaria parasites, is an essential gene family in host-parasite interactions, playing a role in merozoite invasion, parasitophorous vacuole formation, and in the uptake of ions and nutrients from the host plasma^20,21^. The pseudogenization of the CLAG gene on chromosome 8 for *P. vivax-like* suggests that this species lost the CLAG gene during its adaptation to the ape host. The SERA family is needed in the mosquito stages for sporozoites egress from oocysts^22^. In this family, one copy is pseudogeneized in *P. vivax* PvP01 (PvP01_0417400) (Supplementary Figure 4), suggesting that *P. vivax* lost this copy during its evolution in the Anopheles specimen infecting humans.

**Figure 1.**
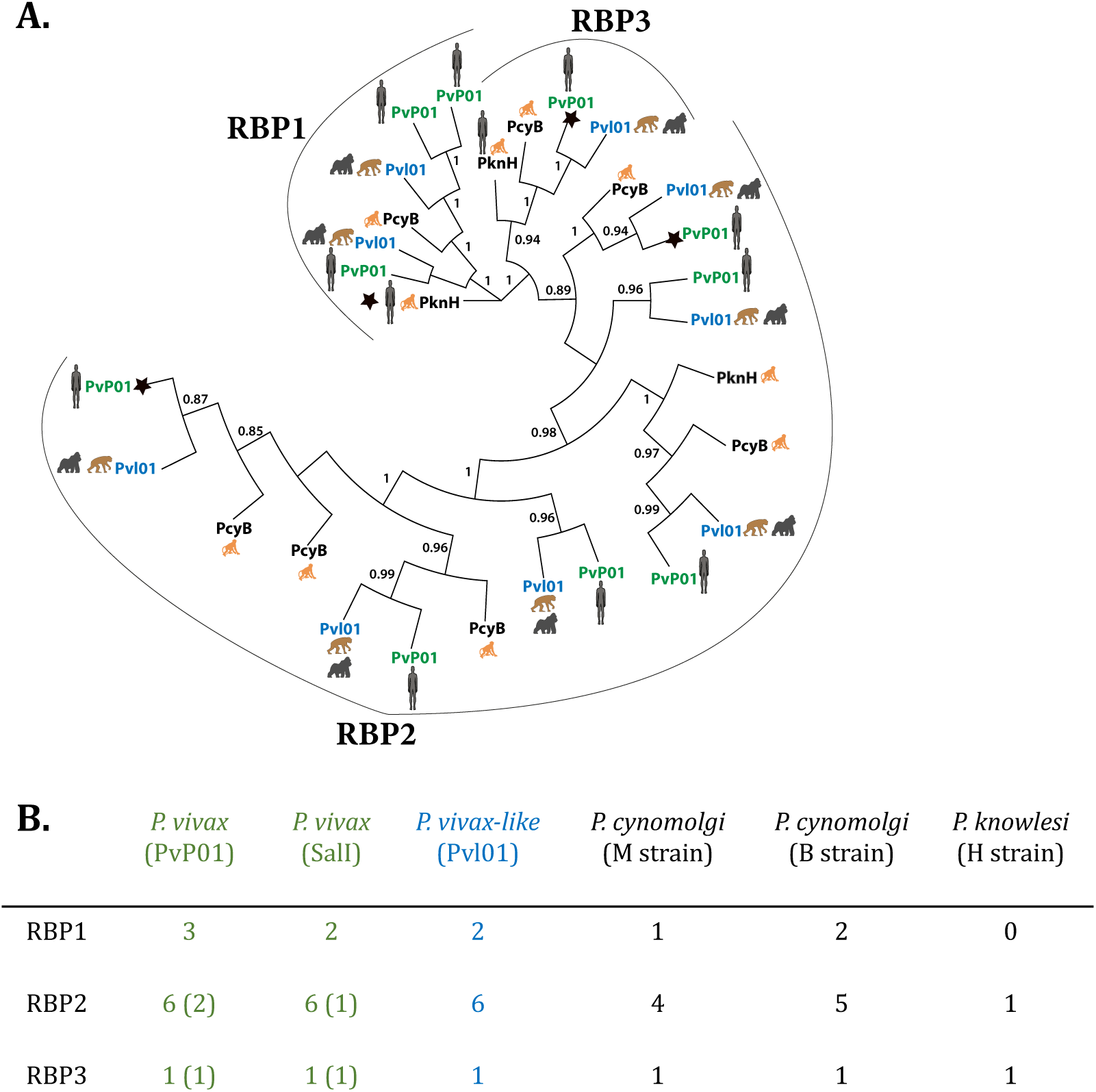
Reticulocyte binding proteins in *P. vivax-like* and *P. vivax*. A. Maximum likelihood phylogenetic tree of all full-length RBPs in *P. vivax-like* Pvl01 (in blue), *P. vivax* SalI and PvP01 strains (in green), P. cynomolgi B strain and P. knowlesi H strain (in black). Only bootstrap values, calculated by RAxML bootstrapping posterior probability, superior to 70% are indicated. The different subclasses of RBPs are indicated as RBP1, RBP2 and RPB3. The stars indicate pseudogenes. The animal pictograms indicate the primate host. B. Table representing the number of variants (including the ones that are pseudogeneized) observed in each RBP subclass in *P. vivax-like* (Pvl01), *P. vivax* (SalI and PvP01), *P. cynomolgi* (B and M strains) and *P. knowlesi* (H strain). Pseudogenes detected among each subclass of RBP are indicated within each subclass between brackets.

**Figure 2.**
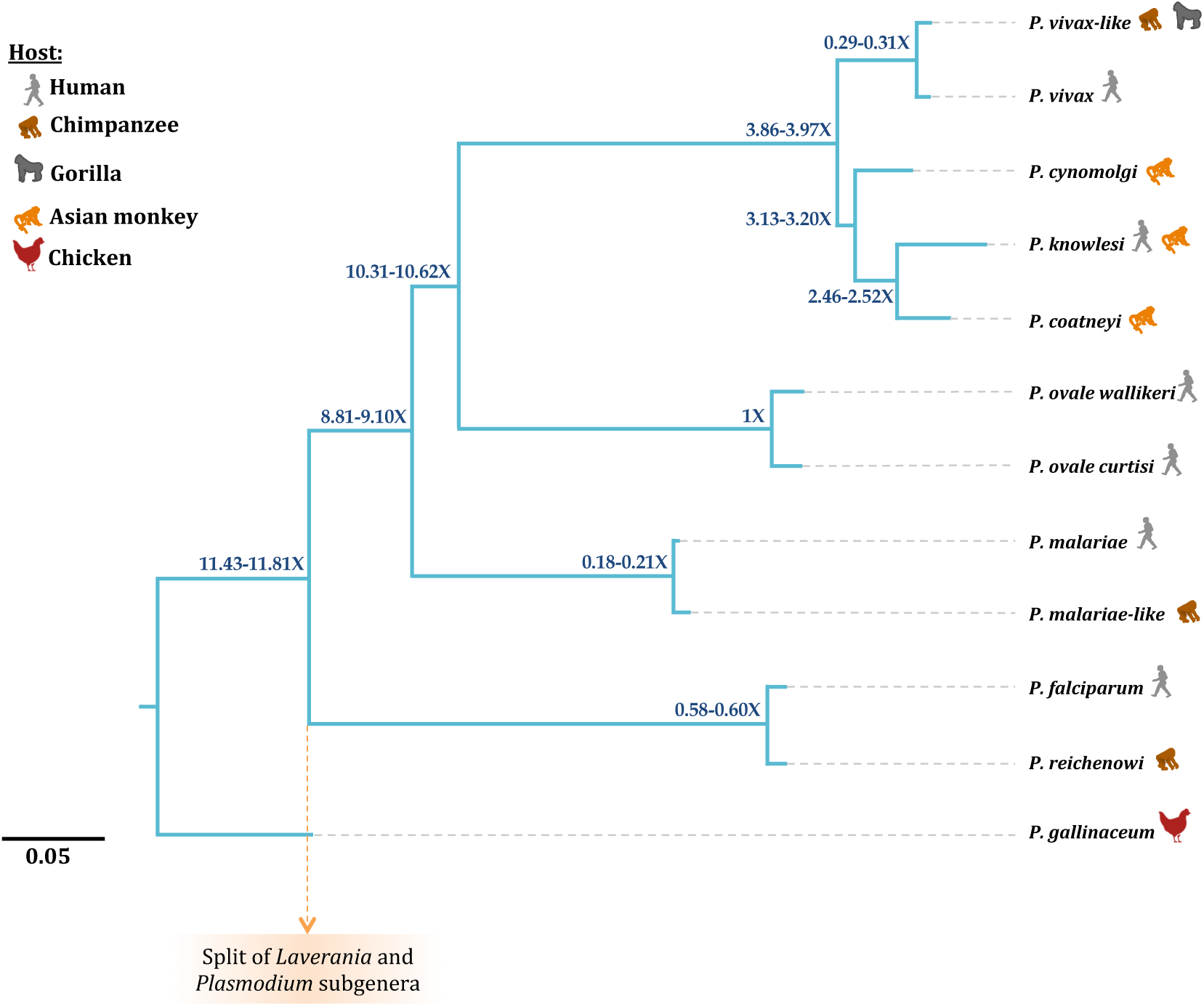
Divergence dating between *P. vivax* and *P. vivax-like*. Maximum likelihood phylogenetic tree of 12 *Plasmodium* species including *P. vivax* and *P. vivax-like*. The analysis was based on an alignment of 2943 1-1 orthologous of 12 *Plasmodium* reference genomes. The relative split between *P. vivax* and *P. vivax-like* is estimated at around three times shorter than the split between *P. ovale wallikeri* and *P. ovale curtisi* and 1.5 times longer than *Plasmodium malariae* and *Plasmodium malariae-like*. The ‘X’ indicates relative split values, based on the fixed point as the relative distance of the split of the speciation of the two *P. ovale* genomes and the *P. malariae-like* and *P. malariae*^14^.

During the life cycle of *Plasmodium* parasites, host RBC invasion is mediated by specific interactions between parasite ligands and host erythrocyte receptors. Two major multigene families are involved in RBC invasion: the Duffy-binding proteins (DBP) and the reticulocyte-binding proteins (RBP)^23^. DBP is a protein secreted by the micronemes of the merozoite stage that binds to the Duffy Antigen Receptor for Chemokine (DARC) to invade RBCs. *P. vivax* is characterized in its genome by two DBP genes (DBP1 on chromosome 6 and DBP2 on chromosome 1) that seem to be essential to RBC invasion, as demonstrated by their inability to infect individuals not expressing the Duffy receptor on the surface of their RBCs (i.e. Duffy-negative individuals)^24–26^. In the reference genomes of *P. vivax-like* (Pvl01 and Pvl06), we observe the DBP1 gene as for the *P. vivax* genome PvP01 (Table 2) and also in the other *Plasmodium* species (*P. knowlesi* and *P. cynomologi*), however we did not observe the DBP2 one (no read obtained mapping to this region). This observation was confirmed in the other genotypes sequenced in this study. Knowing that gorillas and chimpanzees are today all described as Duffy positive^10^, we propose that *P. vivax-like* parasites infect only Duffy positive hosts which could be associated to the absence of the DBP2 gene. This would be in accordance with the fact that the only described transfer of *P. vivax-like* to humans was in a Caucasian Duffy positive individual^9^ and that no transfers of *P. vivax*-*like* were recorded in Central African Duffy negative populations despite the fact that they live in close proximity with infected ape populations^27^.

RBP genes encode a merozoite surface protein family present across all *Plasmodium* species and known to be involved in RBC invasion and host specificity^23^. Among this RBP family genes, three gene classes (RBP1, RBP2 and RBP3) exist and are associated to the ability of *Plasmodium* parasites to invade different maturation stages of RBCs. In this study, comparison of the organization and characteristics of the RBP gene family between *P. vivax, P. vivax-like, P. knowlesi* and *P. cynomolgi* (Figure 1 and Table 2), first reveals that gene classes RBP2 and RBP3 are ancestral to the divergence of all these species except *P. knowlesi*. Then, an expansion of RBP2 class is observed in *P. vivax*/*P. vivax-like/P. cynomolgi* lineage (Figure 1A), suggesting that in this lineage specific expansion likely occurred during the evolution of these species. Finally, RBP3 genes, which are supposed to confer the ability to infect specifically normocytes, are functional in all species except in *P. vivax* (where the gene is pseudogenised in both SalI and PvP01 strains), suggesting that *P. vivax* specifically lost the ability to infect normocytes or have develop an ability to infect specifically only reticulocytes during its adaptation to human’s RBCs (Figures 1A and 1B).

### Phylogenetic relationships to other *Plasmodium* species and divergence time

Conservation of the gene content between *P. vivax-like* with the other primate-infective *Plasmodium* species has enabled us to reconstruct with confidence the relationships between the different species and to estimate the relative age of the different speciation events. This analysis confirmed the position of *P. vivax-like* as the closest sister lineage of *P. vivax* (Figure 2).

Regarding the estimation of divergence times using genomic information, different methods were recently used for *Plasmodium*, such as the one implemented in G-PhoCS^28^ or the one developed by Silva et al^29^. G-PhoCS uses a Bayesian MCMC approach to infer, based on the information provided by multiple loci, the divergence time between species. This method has been applied in two recent studies for *Plasmodium* parasites: one aiming at estimating the relative split times between the two *P. ovale* sub-species and between *P. malariae* and *P. malariae-like*^14^; the other to estimate the divergence time within the *Laverania* subgenus, a subgenus including *P. falciparum* and all its closest ape-relatives^13^. The Silva method is based on the estimate of the sequence divergence in different proteins and comparison of this divergence measured between different lineages^29^. In this method, the regression slope of the divergence between the proteins in two lineages reflects their relative age. The advantage is that it does not relate on an estimate of mutation rate. Finally, it has already been used in a recent study estimating avian and primate *Plasmodium* species divergence times^30^. In this study, we used the Silva method to estimate the divergence time between *P. vivax* and *P. vivax-like* relatively to the other divergence events within the tree. We observe that the evolution of *P. vivax* in humans did not occur at the same time as the other human malaria agents of the *Plasmodium* subgenus (i.e. *P. malariae* and *P. ovale*). Indeed, we show that the time of the split between *P. vivax* and *P. vivax-like* happened before the divergence between *P. malariae* and *P. malariae-like* (about 1.5 times earlier) which itself occurred earlier than the divergence between *P. ovale curtisi* and *P. ovale wallikeri.* The relative divergence times estimated between *P. malariae* / *P. malariae-like* on one side and *P. ovale curtisi* and *P. ovale wallikeri* on the other side are congruent with those obtained previously using the GPhocs method^13,14^. What differs, however, is the relative dating of the split between *P. reichenowi* (a chimpanzee parasite) and *P. falciparum*, two species of the *Laverania* subgenus. While this split is here estimated to have happened earlier than the split between *P. malariae* and *P. malariae-like* (note that similar estimates were recently obtained using another set of data using the Silva method), it was estimated to have occurred at the same time using the GPHOCs method. This discrepancy between methods suggests that the same strict molecular clock (which is a hypothesis of the Silva method) may not apply over the entire tree (especially for the *Laverania* subgenus because of the extreme low GC content of the parasites species of this subgenus in comparison to other *Plasmodium* species). Whatever the method used, all these estimates suggest nevertheless that the evolution of *P. vivax* in humans did not occur at the same time as the other human malaria agents and that the transfer of *Plasmodium* parasites to humans may have happened several times independently over the history of the *Homo* genus.

### Relationships to worldwide human *P. vivax* isolates

To analyse the relationship between our 11 *P. vivax-like* isolates and human *P. vivax*, we completed our dataset with 19 published human *P. vivax* genomes^31^ (Supplementary Table 1). All sequencing reads were aligned against the PvP01 reference genome^15^ and SNPs were called and filtered as described in the Materials and Methods section. Maximum-likelihood phylogenetic trees were then produced based on 100,616 SNPs. Our results clearly demonstrate the presence of a significantly distinct clade (a bootstrap value of 100) composed of *P. vivax-like* strains on one side and human *P. vivax* isolates on the other side (Figure 3). This result differs from previous results suggesting that human strains formed a monophyletic clade within the radiation of ape *P. vivax-like* parasites^10^.

**Figure 3.**
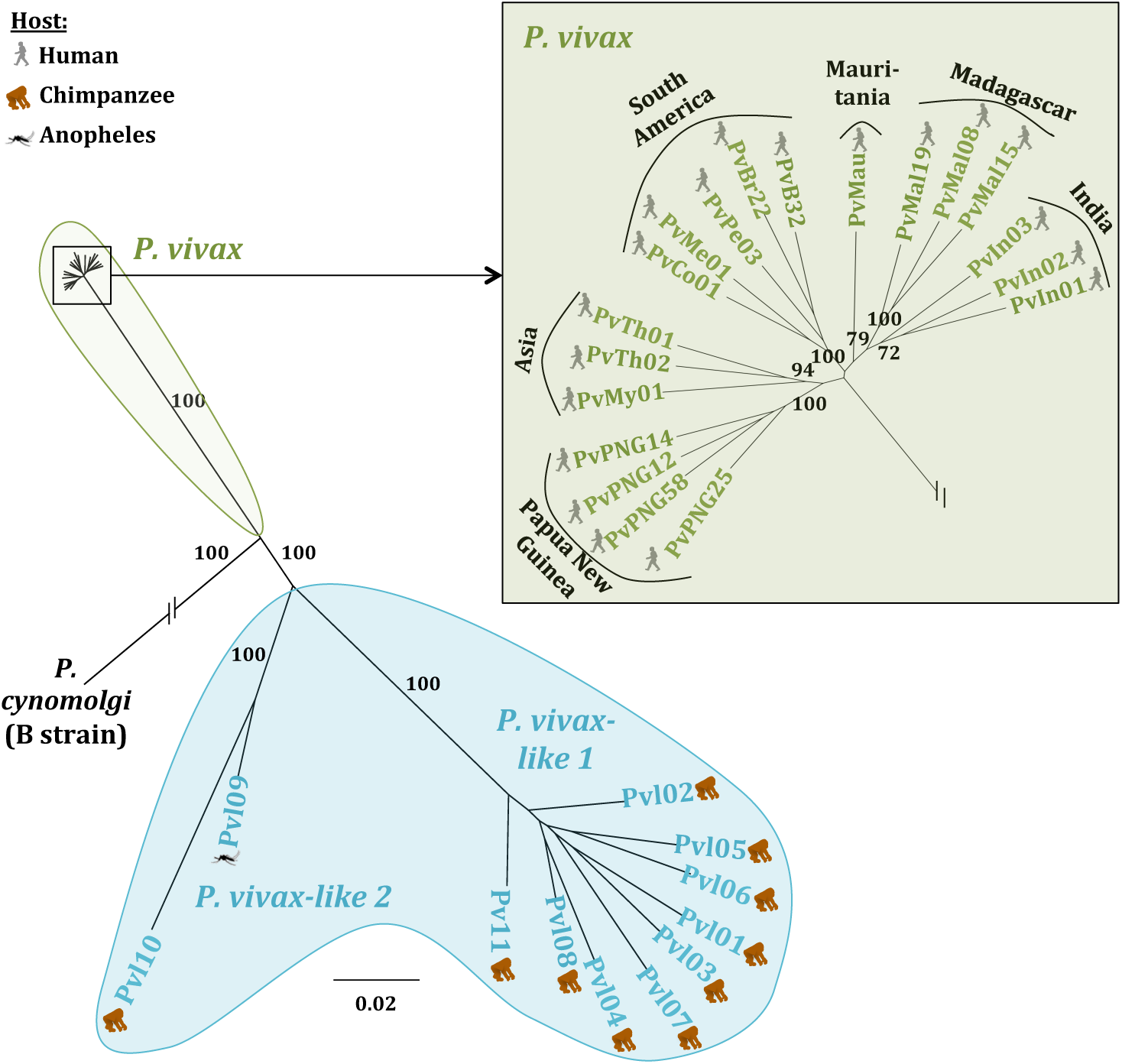
**Maximum likelihood phylogenetic tree** with 1000 bootstraps computed through alignment to the *P. cynomolgi* B strain genome, based on 100,616 SNPs shared by 11 *P. vivax-like* and 19 *P. vivax* samples. Bootstrap values superior to 70% are indicated. The host in which the *Plasmodium* parasite was detected is indicated by the pictograms (human, chimpanzee and *Anopheles*). This phylogeny showed the presence of a significantly distinct clade (high bootstrap values associated to each clade) composed of *P. vivax-like* strains on one side (light blue) and human *P. vivax* isolates on the other side (light green).

One explanation for this difference with previous results could be due to a phenomenon called Incomplete Lineage Sorting (ILS). ILS is the discordance observed between some gene trees and the species or population tree due to the coalescence of gene copies in an ancestral species or population^32^. Such a phenomenon is often observed when species or population divergence is recent, which is the case for *P. vivax*/*P. vivax-like*^33,34^. ILS may thus result in the wrong conclusion of *P. vivax* and *P. vivax-like* populations being intermixed and *P. vivax* diversity being included in the diversity of *P. vivax-like*. In our study, the use of significantly more genetic information from throughout the genome, both in genic and intergenic regions, allows us to reduce this effect of ILS and reflects a more accurate picture of the genetic relationship between the different parasite species. Clearly for now, the origin, the direction of the transfer and the evolutionary history of these parasites are still unclear and need the addition of more *P. vivax-like* samples from other locations to be included.

Our results also show that *P. vivax-like* is composed of two distinct lineages: the one including the two reference genomes (Pvl01 and Pvl06) and seven other isolates that will hereafter be referred as *P. vivax-like 1* and another one including two isolates (Pvl09 and Pvl10) (referred as *P. vivax-like 2*) (Figure 3). These two lineages may reflect an ancient split within *P. vivax-like* or be the consequence of a recent introgression or hybridization event between *P. vivax-like* and *P. vivax* in Africa. Further analyses including sequencing of more African *P. vivax* populations from different geographic areas, should be done to disentangle these two hypotheses.

Previous studies highlighted the high genetic diversity of *P. vivax-like* populations in comparison to *P. vivax* worldwide^15^. In this genome-wide analysis of the nucleotide diversity π^35^, we confirm that *P. vivax-like* populations are significantly more diverse than *P. vivax* populations (P<0.001, Wilcoxon test), with *P. vivax-like* samples showing nearly ten times higher nucleotide diversity (π_*P.vivax*_ = 0.0012; π_*P.vivax-like*_ = 0.0096). This suggests that African great apes *P. vivax-like* parasites are probably more ancient that the human *P. vivax* strains and that the human *P. vivax* species (as other human *Plasmodium* species) went through a bottleneck and only recently underwent population expansion.

### *P. vivax* specific adaptive evolution

Comparison of the *P. vivax* genome to its closest sister lineage (*P. vivax-like*) and to the other primate *Plasmodium* provides a unique opportunity to identify *P. vivax* specific adaptations to humans. We applied a branch-site test of positive selection to detect events of positive selection that exclusively occurred in the *P. vivax* lineage. Within the reference genome *P. vivax-like* (Pvl01), 418 genes exhibited significant signals of positive selection (Supplementary Table 3). In the human *P. vivax* genome PvP01, the test allowed the identification of 255 genes showing significant signals of positive selection (Supplementary Table 4). Among these genes presenting a significant dN/dS ratio, 71 were shared between *P. vivax* and *P. vivax-like*, including 56 encoding for proteins with unknown function, and 15 encoding for proteins that are involved either in energy metabolism regulation (n = 9), in chromatid segregation (n = 2) or cellular-based movement (n = 4).

We then took into consideration the genes detected under positive selection in *P. falciparum*^13^ and compared them to those obtain in *P. vivax*. We identified of a subset of 10 genes under positive selection in the human *P. vivax* and *P. falciparum* parasites (P-value<0.05) (Supplementary Table 5). Among these 10 genes, five are coding for conserved *Plasmodium* proteins with unknown function and three for proteins involved in either transcription or transduction. Interestingly, the two remaining genes under positive selection in these two human *Plasmodium* parasites code for the oocysts capsule protein, which is essential for malaria parasite survival in *Anopheles*’ midgut, and for the rhoptry protein ROP14, involved in the protein maturation and the host cell invasion. These results suggest that these proteins could be essential for infection of humans or their vectors and future studies should focus on the involvement of these proteins in human parasite transmission and infection.

## Conclusion

Through technical accomplishments we produced and assembled the first *P. vivax-like* reference genomes, the closest sister clade to human *P. vivax*, which is an indispensable step in the development of a model system for a better understanding of this enigmatic species. We established that *P. vivax-like* parasites form a genetically distinct clade from *P. vivax*. Concerning the relative divergence dating, we estimated that the divergence between both species occurred probably before the split between *Plasmodium malariae*. This suggests that the transfer of *Plasmodium* parasites to humans happened several times independently over the history of the *Homo* genus. Our genome-wide analyses provided new insights into the adaptive evolution of *P. vivax*. Indeed, we identified several key genes that exhibit signatures of positive selection exclusively in the human *P. vivax* parasites, and show that some gene families important for RBC invasion have undergone species-specific evolution in the human parasite, such as for instance RBPs and DBPs. Are these genes the keys of the emergence of *P. vivax* in the human populations? This pending question will need to be answered through functional studies associated to deeper whole genome analyses. Interestingly, among the genes identified under positive selection, two have been identified to also be under positive selection in the other main human malaria agent, *P. falciparum*, thus suggesting their key role in the evolution of the ability of these parasites to infect humans or their anthropophilic vectors. To conclude, this study provides the foundation for further investigations into *Plasmodium vivax* parasite’s traits of public health importance, such as features involved in host-parasite interactions, host specificity, and species-specific adaptations.

## Material and methods

### *P. vivax-like* sample collection and preparation

*P. vivax-like* samples were identified by molecular diagnostic testing during a continuous survey of great ape *Plasmodium* infections carried out in the Park of La Lékédi, in Gabon, by the Centre International de Recherches Médicales de Franceville (CIRMF)^9^. In parallel, a survey of *Anopheles* mosquitoes circulating in the same area (Park of La Lékédi, Gabon) was conducted in order to identify potential vectors of ape *Plasmodium*^11^. Specifically, mosquitoes were trapped with CDC light traps in the forest of the Park of La Lékédi in Gabon. *Anopheles* specimens were retrieved and identified using a taxonomic key^36^ before proceeding to dissection to isolate the abdomen. Samples were then stored at -20°C until transportation to the CIRMF, Gabon, where they were stored at -80 °C until processed. Blood samples of great apes were treated using leukocyte depletion by CF11 cellulose column filtration^37^. *P. vivax-like* samples were identified either by amplifying and sequencing the *Plasmodium Cytochrome b* (*Cytb*) gene as described in Ollomo et al. or directly from samples already studied for other *Plasmodium* species^29,38^. This allowed the detection of 11 *P. vivax-like* samples, 10 from chimpanzees and 1 from an *Anopheles moucheti* mosquito. Most of these samples were co-infected with other *Plasmodium* species, and/or probably with multiple *P. vivax-like* isolates (see below and Supplementary Table 1). The identification of intraspecific *P. vivax-like* co-infections was made by analyzing the distribution of the reference allele frequency^38^.

### Ethical approval

These investigations were approved by the Government of the Republic of Gabon and by the Animal Life Administration of Libreville, Gabon (no. CITES 00956). All animal work was conducted according to relevant national and international guidelines.

### Genome sequencing

DNA was extracted using Qiagen Midi extraction kits (Qiagen) following manufacturer’s recommendations, and then enriched through a whole genome amplification step (WGA^39^). The Illumina isolates were sequenced using Illumina Standard libraries of 200-300bp fragments and amplification-free libraries of 400-600bp fragments were prepared and sequenced on the Illumina HiSeq 2500 and the MiSeq v2 according to the manufacturer’s standard protocol (Supplementary Table 1). The Pvl06 isolate was sequenced using Pacific Biosciences with the C3/P5 chemistry after a size selection of 8 kb fragments. Raw sequence data are deposited in the European Nucleotide Archive. The accession numbers can be found in Supplementary Table 1.

### Assembly of *P. vivax-like* genomes

Two *P. vivax-like* genomes (Pvl01 and Pvl06) were assembled from a co-infection with a *P. malariae-like* and a *P. reichenowi* (PmlGA01 sample in Rutledge et al. 2017)^14^ for Pvl01 and from a co-infection with *P. gaboni* for Pvl06 (PGABG01 sample in Otto et al.)^40^. Briefly, the genome assembly of the Illumina sequenced sample Pvl01 was performed using MaSuRCA^41^ and the assembled contigs belonging to *P. vivax-like* were extracted using a BLAST search against the *P. vivax* P01 reference genome (PvP01 genome; http://www.genedb.org/Homepage/PvivaxP01)^15^. The draft assembly was further improved by iterative uses of SSPACE^42^, GapFiller^43^ and IMAGE^44^. The 3,540 contigs resulting from these analyses were then ordered against the PvP01 genome and the *P. gaboni* and *P. reichenowi* reference genomes^13^ to separate possible co-infections with a parasite species of chimpanzees from the *Laverania* subgenus using ABACAS2^45^. The genome assembly was further improved and annotated using the Companion web server^16^. BLAST searches of the unassembled contigs against the two reference genomes were performed before running Companion to keep the contigs with the best BLAST hits against PvP01 only. The PacBio assembly of Pvl06 was performed using the Hierarchical Genome Assembly Process HGAP^46^.

### Read mapping and alignment

Nine additional *P. vivax-like* samples were sequenced for population genomics and polymorphism analyses (see Supplementary Table 1). The dataset was completed with 19 globally sampled *P. vivax* isolates^31^ for human *vs.* great apes parasite comparisons, and the Asian parasite *P. cynomolgi* strain B was used as the root for phylogenetic inferences^8^. The 11 newly generated *P. vivax-like* samples, together with the already published 19 *P. vivax* samples and the reference strain *P. cynomolgi*^8^ Illumina reads were mapped against the PvP01 reference genome using BWA^47^ with default parameters. We then used Samtools to keep only the properly paired reads and to remove PCR duplicates^48^.

### Gene family search

For all *P. vivax-like* Pvl01 and Pvl06, *P. vivax* PvP01 and SalI, *P. cynomolgi* B strain and *P. knowlesi* H strain genomes obtained, gene variants were detected and counted using Geneious software^49^.

### Orthologous group determination and alignment

Orthologous groups across (1) *P. vivax* PvP01, *P. vivax-like* Pvl01, *P. cynomolgi* B strain^8^, *P. cynomolgi* M strain^50^ and *P. knowlesi* H strain^17^ reference genomes and (2) the 13 *Plasmodium* reference genomes used for the phylogeny (the seven *Laverania* genomes *P. falciparum*^51^, *P. praefalciparum, P. reichenowi, P. billcollinsi, P. blacklocki, P. gaboni* and *P. adleri*^13^, *P. cynomolgi* B strain and *P. knowlesi* H, *P. vivax* PvP01, *P. vivax-like* Pvl01 (this study), and *P. malariae* and *P. malariae-like* ^14^ were identified using OrthoMCL v2.09^52,53^. From those, we extracted different sets of one-to-one orthologues for the subsequent analyses: a set of 4,056 genes that included the one-to-one orthologues among the four restricted species, *P. vivax, P. vivax-like, P. cynomolgi* and *P. knowlesi*, and a set of 2,352 among the 13 *Plasmodium* species considered here for the interspecies phylogenetic analysis.

Amino acid sequences of the one-to-one orthologues were aligned using MUSCLE^54^. Prior to aligning codon sequences, we removed the low complexity regions identified on the nucleotide level using dustmasker^55^ and then in amino acid sequences using segmasker ^56^ from ncbi-blast. After MUSCLE alignments^54^, we finally excluded poorly aligned codon regions using Gblocks default parameters^57^.

### SNP discovery and annotation

SNPs were called independently for all 11 *P. vivax-like* and 19 *P. vivax* samples by first mapping the samples against the *P. vivax* PvP01 reference genome using SMALT and then calling SNPs using Samtools mpileup v. 0.1.9 (parameters –q 20 -Q 20 -C 50) followed by bcftools (call -c -V indels). SNPs were filtered using VCFTools (--minDP 5 –max-missing 1).

### Divergence dating

To estimate the dates of speciation, we used 12 *Plasmodium* genomes: the here generated *P. vivax-like* Pvl01, *P. vivax* PvP01^15^, *P. cynomolgi* M Version 2^50^, *P. coatneyi* PcyM^50^, *P. knowlesi* H strain^17^, *P. falciparum* 3D7^51^ *P. reichenowi* PrCDC^40^, *P. gallinaceum*^58^, and *P. ovale wallikeri, P. ovale curtisi, P. malariae* and *P. malariae-like*^14^. From the proteins of the 12 genomes, low complexity regions were excluded with SEG filter, using default parameters^59^. After an all-against-all BLASTp (parameter Evalue 1e-6), OrthoMCL v.1.4^53^ (using default parameters) was run. For each of the 2943 1-1 orthologous, an alignment was generated with MUSCLE^54^ and the alignment was finally cleaned with Gblocks (parameters: -t=p -b5=h -p=n -b4=2)^60^.

To build the phylogenetic tree, the software RAxML v.8.2.8^61^ was used on the concatenated alignments of 1000 random picked orthologous. The PROTGAMALG substitution model was then used, as proposed in Rutledge et al^14^, 100 bootstraps were run confirming the tree.

To date the speciation events, the method from Silva et al.^29^ was applied. The dAA was obtained through a pairwise comparison using PAML v.4.7 ^62^. An R script from the authors of the method^29^ allowed the estimation of alpha with the error bound for each pair, based on a Total Least Squares regression. Results are reported in Figure 2. As a fix point, we used the relative distance of the split of the speciation of the two *P. ovale* genomes and the *P. malariae-like* and *P. malariae*^14^. The split of *P. reichenowi* and *P. falciparum* was also dated based on the *P. malariae* and *P. malariae-like* split estimation^14^. However, this will need to be confirmed with other methods, because the GC content could bias this estimation. Indeed, the models used in our study assume a strict molecular clock, which would not apply to all *Plasmodium* species, specifically for *P. falciparum* because of its extreme GC content in comparison to other *Plasmodium* species.

### Phylogenetic tree of *P. vivax* and *P. vivax-like* strains

For Figure 1, we performed a maximum likelihood analysis using RAxML^61^ with 100 bootstrap replicates. The tree was visualized using Geneious software^49^. For Figure 3, we constructed a maximum-likelihood tree using the filtered variant call set of SNPs limited to the higher allelic frequency genotypes identified within each sample using RAxML and PhyML (using general-time reversible GTR models)^61,63^. Trees were visualized using Geneious software^49^. All approaches showed the same final phylogenetic tree described in the results section.

### Genome wide nucleotide diversity

For the *P. vivax* and *P. vivax-like* populations, we calculated the genome-wide nucleotide diversity (π)^35^ using VCFTools^64^. The nucleotide diversity was compared between *P. vivax* and *P. vivax-like* species based on the Wilcoxon-Mann-Whitney non-parametric test.

### Detection of genes under selection

In order to identify genomic regions involved in parasite adaption to the human host, i.e. regions under positive selection, we performed branch site tests. To search for genes that have been subjected to positive selection in the *P. vivax* lineage alone, after the divergence from *P. vivax-like*, we used the updated Branch-site test of positive selection^65^ implemented in the package PAML v4.4c^62^. This test detects sites that have undergone positive selection in a specific branch of the phylogenetic tree (foreground branch). All coding sequences in the core genome were used for the test (4,056 gene sets of orthologous genes). A set of 4056 orthologous groups between *P. vivax, P. vivax-like, P. knowlesi* and *P. cynomolgi* was used for this test. dN/dS ratio estimates per branch and gene were obtained using Codeml (PAML v4.4c) with a *free-ratio* model of evolution^62^.

### Data availability

All sequences are being submitted to the European Nucleotide Archive. The accession numbers of the raw reads and assembly data will be found in Supplementary Table 2. As the assemblies are private, they will be available on request.

## Acknowledgements

Authors thank ANR ORIGIN JCJC 2012, LMI ZOFAC, CNRS-INEE, CIRMF, IRD, Sanger Institute for financial support and Société d’Exploitation du Parc de la Lékédi, Bakoumba, GABON. GGR is supported by the Medical Research Council (MR/J004111/1) and the Wellcome Trust (098051). Authors thank D. Green for her comments on this work.

## Author contributions

DTO, FR, FP and VR designed the study. CA, PD, BO, NDM, APO, BN, BM, LB, CP, FP and VR collected and assessed samples. CA performed the WGA. TDO managed the sequencing. AG, BF and TDO did assembly and annotation. TDO, VR and FP performed the evolutionary analyses on core genomes. TDO and GGR performed the dating analyses. AG, TDO, FR and VR wrote the manuscript. All authors read and approved the paper.

## Materials & Correspondance

Rougeron Virginie; Laboratoire MIVEGEC (UM-CNRS-IRD), 34394 Montpellier, France;

